# Whole-transcriptome analysis of BLV-infected cows reveals downregulation of immune response genes in high proviral loads cows

**DOI:** 10.1101/2024.12.23.627929

**Authors:** MI Petersen, G Suarez Archilla, MM Miretti, KG Trono, HA Carignano

## Abstract

Bovine leukemia virus (BLV) is a retrovirus that infects cattle, causing bovine enzootic leukosis, a chronic disease characterized by the proliferation of infected B cells. BLV proviral load (PVL) is a key determinant of disease progression and transmission risk. Cattle can exhibit distinct phenotypes of low PVL (LPVL) or high PVL (HPVL), which remain stable throughout their lifetime. Differential expression analysis revealed 1,908 differentially expressed genes (DEGs) between HPVL and LPVL animals, including 774 downregulated (DReg) and 1134 upregulated (UReg) genes. Functional enrichment analysis revealed that DReg genes were associated primarily with immune response pathways. Conversely, the UReg genes were enriched in processes related to cell cycle regulation, mitotic division, and DNA biosynthesis. Protein□protein interaction analysis revealed six highly interconnected clusters. Interestingly, a cluster was enriched for sphingolipid metabolism, a process critical to enveloped virus infection and immune receptor signaling. These findings provide valuable insights into the molecular mechanisms of BLV infection, suggesting potential markers for disease monitoring and targets for therapeutic intervention.

## Introduction

Bovine leukemia virus (BLV) is a retrovirus that naturally infects cattle, causing a chronic disease known as bovine enzootic leukosis (Bendixen, 1963). BLV primarily infects mature B cells (Meirom et al., 1997), although other blood cells can also be affected (Schwartz and Lévy, 1994; Panei et al., 2013).

After infection, the viral RNA is retrotranscribed and integrated randomly into the host cell genome as a provirus. An increase in the blood proviral load (PVL) is correlated with disease progression (Mirsky et al., 1996; Jimba et al., 2010; Panei et al., 2013; Sato et al., 2018) and the risk of BLV transmission to a healthy herd mate (Gutiérrez et al., 2014b; Juliarena et al., 2016).

In dairy cattle populations, two distinct groups of animals can be distinguished: low PVL (LPVL) and high PVL (HPVL) and, generally, the PVL level remains stable throughout an individual cow’s lifetime (Gutiérrez et al., 2014b; Merlini et al., 2016; Hutchinson et al., 2021).

Several strategies have been proposed for controlling BLV dissemination, including the use of BLV attenuated vaccines (Suárez Archilla et al., 2022) and culling HPVL cows from herds (Ruggiero et al., 2019). Alternatively, genetic polymorphisms in the major histocompatibility complex class II *BoLA-DRB3* gene have been suggested as markers for selecting and breeding LPVL cows (Mirsky et al., 1998; Juliarena et al., 2008; Jimba et al., 2012; Miyasaka et al., 2013; Carignano et al., 2017; Takeshima et al., 2019). However, the genetic basis of disease resistance/susceptibility is complex and polygenic, influenced by multiple loci with small effects on the phenotype (Minozzi et al., 2012; Sahana et al., 2013; Neibergs et al., 2014; Brym and Kamiński, 2017; Carignano et al., 2018).

Genomic association studies have revealed that single-nucleotide polymorphisms (SNPs) linked to BLV PVL variation in infected cattle are distributed across the entire bovine genome (Takeshima et al., 2017; Carignano et al., 2018). In a previous work, we evaluated the expression of candidate genes located within regions harboring associated SNPs in HPVL and LPVL cows. Our results revealed differential expression of the *ABT1* transcription factor and a significant correlation between lymphocyte count and *PRRC2A* and *IER3* gene expression. These genes had not previously been associated with BLV infection phenotypes (Petersen et al., 2021).

Few investigations assessed the global gene expression response to BLV (Bai et al., 2020), and compared naturally BLV-infected vs. noninfected animals (Brym and Kamiński, 2017; Ashrafi et al., 2020; Nishimori et al., 2023). Depending on the BLV phenotype evaluated and/or the route of infection considered, differentially expressed genes have been found to be related to DNA mismatch repair, cell cycle and growth factors (Ashrafi et al., 2020; Bai et al., 2020; Nishimori et al., 2023), as well as innate and adaptive immunity, including complement system activation (Brym and Kamiński, 2017). Nishimori et al. (2023) compared healthy animals, BLV-infected animals, and BLV-infected animals with lymphoma and identified host gene expression correlated with the BLV proviral load, potential markers for monitoring disease progression.

However, the mechanisms underlying BLV PVL development and progression in cattle remain poorly understood. In the present study, we assessed the whole-transcriptome response of BLV-infected cows exhibiting contrasting PVL levels to identify the host genes and metabolic pathways involved. Differential expression analysis allowed us to identify a group of significantly differentially expressed genes (DEGs) when comparing HPVL and LPVL cows. These DEGs were functionally annotated into biological categories, providing insight into the underlying mechanisms of BLV infection.

## Materials and methods

### Sample selection and collection

BLV-infected cows were selected from previously phenotyped animals (Petersen et al., 2021). Briefly, 129 adult Holstein cows from the Argentinian main dairy farm region (31°16′S, 61°29′W) were sampled. Animals, which shared the same lactation period were screened via anti-BLV ELISA (see below) twice, at −10 mo (T1) and −5 mo (T2) from the final sampling time; the mean percentage of reactivity (PR) was 122.7 ± 34.8 and 146.3 ± 55.6, respectively. Then, individuals in the lowest and highest PR quartiles (q) [T1-q_1_ = 25.0–102.8% and T2-q_1_ = 25.0–118.2% and T1-q_4_ = 148.6–178.6% and T2-q_4_ = 194.4–239.6%, respectively] at both times were selected for proviral load (PVL) determination. The PVL phenotype was measured by qPCR (see below) in 15 cows twice, at −3 mo (T3) and 0 mo (T4). Finally, 6 cows with consistent high-PVL (HPVL) and 6 with low-PVL (LPVL) were selected.

Cow fresh blood samples (45 ml) were obtained via jugular venipuncture and supplemented with EDTA (225 μM). Peripheral blood mononuclear cells (PBMCs) were isolated via Ficoll□Paque Plus (GE Healthcare, Uppsala, Sweden) density gradient centrifugation (following the manufacturer’s protocol) and resuspended in RNAlater® (Ambion, Austin, TX, USA). Sample collection and PBMC isolation were performed on the same working day to prevent any RNA degradation. The samples were stored at -80°C until use.

The animal handling and sampling procedures followed the manual’s recommendations of the Animal Care and Use Committee of the National Institute of Agricultural Technology (INTA, Buenos Aires, Argentina).

### Anti-BLV ELISA

An indirect ELISA against the whole BLV viral particle-lysed antigen was used following a previously described protocol (Trono et al., 2001). A weak positive international control standard serum was utilized as a reference to calculate a normalized sample-to-positive ratio. The difference between the optical density values obtained for the weak positive control and negative BLV control samples was set as 100% reactivity. All the tested samples were referred to it. Samples with PRs above the cutoff level (>25%) were considered positive.

### BLV PVL quantification

Genomic DNA (gDNA) was isolated from PBMCs via a High Pure PCR Template Preparation Kit (Roche, Penzberg, Germany). The concentration (ng/μL) and quality (A260/280 and A260/230) of the gDNA were measured via a spectrophotometer (Nanodrop, Thermo Fisher Scientific, Waltham, MA).

To quantify the BLV PVL, validated SYBR Green dye-based qPCR targeting the BLV *Pol* gene was used (Petersen et al., 2018). Briefly, each qPCR mixture (final volume = 25 μL) contained Fast Start Universal SYBR Green Master Mix (Roche), forward and reverse primers (800 nM; BLVpol_5f: 5′-CCTCAATTCCCTTTAAACTA-3′; BLVpol_3r: 5′-GTACCGGGAAGACTGGATTA-3′; Thermo Fisher Scientific), and 200 ng of gDNA template. The amplification and detection reactions were performed via a Step One Plus device (Applied Biosystems, Foster City, CA).

The thermocycler profile included predenaturation for 2□min at 50°C, then denaturation for 10□min at 95°C, followed by 40 cycles of 15□s at 95°C, 15□s at 55°C, and a final extension of 60□s at 60°C. The specificity of each BLV-positive reaction was confirmed through melting temperature dissociation curve (T_m_) analysis. Samples with PVL values <1,500 copies/μg of total DNA were considered LPVL; otherwise, they were considered HPVL (Petersen et al., 2021).

### RNA isolation and library sequencing

Total RNA was extracted from PBMCs stored in RNAlater® with a High Pure RNA Isolation Kit (Roche) via a modified protocol. The DNase treatment step was set to 20 min at 37°C. The RNA concentration and integrity were evaluated using a 2100 Bioanalyzer chip (Agilent Technologies, Santa Clara, CA, USA) (Figure S1). RNA samples with a RIN value ≥ 8 were used to prepare sequencing libraries with the TruSeq Stranded mRNA Kit (Illumina, San Diego, CA, USA) following the manufacturer’s protocol.

Samples from four animals per phenotypic group were sequenced (HPVL: A1, A2, A6 and A7; LPVL: B2, B5, B6 and B7). Equal molar concentration sequencing libraries were pooled and set in 4 sequencing lanes of a NextSeq500 equipment (Illumina) run to avoid possible batch effects (Li et al., 2014). A total of 354.4 M reads (1 × 75 bp) were obtained.

### Quality control and reference genome mapping

Demultiplexed Fastq files for each sample were visually inspected with FastQC (Andrews, n.d.). Quality control (QC) and adapter trimming were performed with TrimGalore v0.6.4 (Krueger et al., 2021) using a q score >20. Reads with lengths < 20 bp were discarded. QC-passed reads were mapped to the bovine reference genome ARS-UCD1.2 v100 (Rosen et al., 2020) using HISAT2 v2.2.0 (Kim et al., 2019). The mapped read statistics were evaluated with RSeQC v4.0.0 (Wang et al., 2012) (Table S1).

### Gene expression quantification and differential expression analysis

Genomic features (genes) were quantified via HTSeq-count v0.12.4 (Anders et al., 2015) in the “union” mode, which counts only uniquely mapped reads and discards those overlapping with two or more genes. This process yielded a raw count expression matrix for each sample and annotated gene (8 × 27607). The expression matrix was subsequently analyzed using the edgeR package (Robinson et al., 2010; McCarthy et al., 2012) within the Bioconductor project v3.13 (Gentleman et al., 2004). All the analyses were conducted in the R environment.

The relationships between the gene expression profiles of the HPLV and LPVL samples were visualized using a multi-dimensional scaling plot. The leading 500 gene log2-fold changes (root-mean-square average) between each pair of samples were used as distances.

Differentially expressed genes (DEGs) between the HPVL and LPVL groups were tested via an exact test based on the quantile-adjusted conditional maximum likelihood (qCML) implemented in edgeR (Robinson and Smyth, 2008). Multiple comparisons correction of the p values was performed by the Benjamini-Hochberg false discovery rate (BH-FDR) method (Benjamini and Hochberg, 1995). BH-FDR adjusted p values (q-value) < 0.05 were considered statistically significant.

### Functional annotation and protein□protein interaction network analysis

Functional inference of each expressed gene were made using PANTHER database (db) v17.0 (https://www.pantherdb.org/) (Mi et al., 2019, 2021) and STRING db (https://cn.string-db.org/) (Szklarczyk et al., 2011, 2021). The PANTHER ontological classification system provides functional categories integrating experimental evidence, functional and evolutionary information on protein family phylogenetic trees (Thomas et al., 2003; Mi et al., 2019), and the Protein Class (PC) and abbreviated Gene Ontology (GO-Slim) terms were used. In addition, the Kyoto Encyclopedia of Genes and Genomes (KEGG) metabolic pathways (http://www.genome.jp/kegg/) (Kanehisa and Goto, 2000) were retrieved using STRING db.

Enrichment analysis of GO/PC PANTHER/KEGG metabolic pathway terms was performed on the list of 13382 expressed genes (EGs) obtained after filtering out genes whose expression was low. These genes were ranked by their logFC values between the HPVL and LPVL groups. The Mann□Whitney rank-sum test (U test) was used to indicate functional categories that are more overrepresented at the top or bottom (up-or down-regulated genes) of the ranked list than expected by chance. Enrichment test p-values were adjusted for multiple testing with the BH-FDR method, and q-values < 0.05 were considered statistically significant.

A protein□protein interaction (PPI) network of DEGs was constructed via STRING db, which compiles curated known and predicted protein□protein interactions from several sources, i.e., experimental evidence, computational prediction, scientific literature, coexpression, other (primary) databases, etc. The resulting network was visualized in Cytoscape v3.9.0 (Shannon et al., 2003). Clusters of highly interconnected proteins were identified using the molecular complex detection algorithm implemented in MCODE v2.0 with default parameters (Bader and Hogue, 2003). An overrepresentation analysis of GO terms and KEGG metabolic pathways was applied to clusters via Fisher’s exact test on the basis of the hypergeometric distribution (Tavazoie et al., 1999).

### Gene expression by RT□qPCR

A total of 5 statistically significant (FC >1.5) DEGs (*BLNK, PIK3CA, BoLA-DQB, CD8A* and *CD4*) were selected for RT□qPCR validation. The assay was conducted following the recommendations proposed by the minimum information for publication of quantitative real-time PCR experiments following the minimum recommendations (MIQE) guidelines (Bustin et al., 2009). The primers (Table S2) were designed using the Primer-BLAST tool (Ye et al., 2012). Homo and heterodimers, GC content, Tm, and potential secondary structures for each primer pair were evaluated using OligoAnalyzer software v3.1 (Owczarzy et al., 2008).

RNAs (300 ng) extracted from 5 HPVL and 5 LPVL (Table S3, Petersen et al., 2021) were treated with the DNase RQ1 (Promega, US). Then, the MultiScribe High-Capacity cDNA Reverse Transcription Kit (Applied Biosystems, US) was used for cDNA synthesis, according to the manufacturer’s instructions.

RT□qPCR was performed with a 1:10 dilution of cDNA using the SsoAdvanced Universal SYBR Green Supermix (Bio-Rad, US) in a final reaction volume of 10 μL, containing 300 nM of each specific primer (forward or reverse) and 3 μL of cDNA. Runs were performed with StepOne Plus equipment (Applied Biosystems, US) following standard cycling conditions: (1) 95°C for 30 s and (2) 40 cycles of 95°C for 15 s and 60°C for 1 min. Reaction specificities were confirmed by inspection of the T_m_ dissociation curve. Technical duplicates were assayed for all RT□qPCRs. For each gene, non-RT RNAs and no template controls were incorporated. The reaction efficiencies were calculated using the LinRegPCR software (Ramakers et al., 2003). Relative expression values were normalized against two bovine reference genes, *RPLP0* and *B2M* (Brym et al., 2013). Differential expression analysis was performed via REST software (Pfaffl et al., 2002). This returns a relative expression value (R): an R > 1 indicates higher gene expression in the HPVL phenotype than in the LPVL phenotype, whereas an R < 1 indicates lower expression.

## Results

An average mapping rate exceeding 97% was achieved, with more than 85% of the reads uniquely mapped to the bovine reference genome. Additionally, approximately 69% of the reads aligned to exonic regions.

Unsupervised clustering revealed that sample A2 had a distinct expression profile from that of the HPVL group (Figure S2). Removing A2 improved the clustering of LPVL and HPVL samples, so we excluded it from downstream analysis.

The differential expression analysis between HPVL (n=3) and LPVL (n=4) was performed on expressed genes (EGs) identified 2216 differentially expressed genes (DEGs) (q < 0.05) out of 13,382 genes evaluated. Among these, 86.1% (1,908) showed moderate changes (FC ≥ |1.5|), 37.4% (829) of DEGs exhibited high fold change values (FC ≥ |2|), and 0.85% (19) showed differences exceeding a FC >10 (Tablexx). From those 1908 genes with fold changes (FCs) ≥ |1.5|, 774 were downregulated (DReg) and 1134 upregulated (UReg) genes (Figure 1, Table S4).

**Figure 1.**
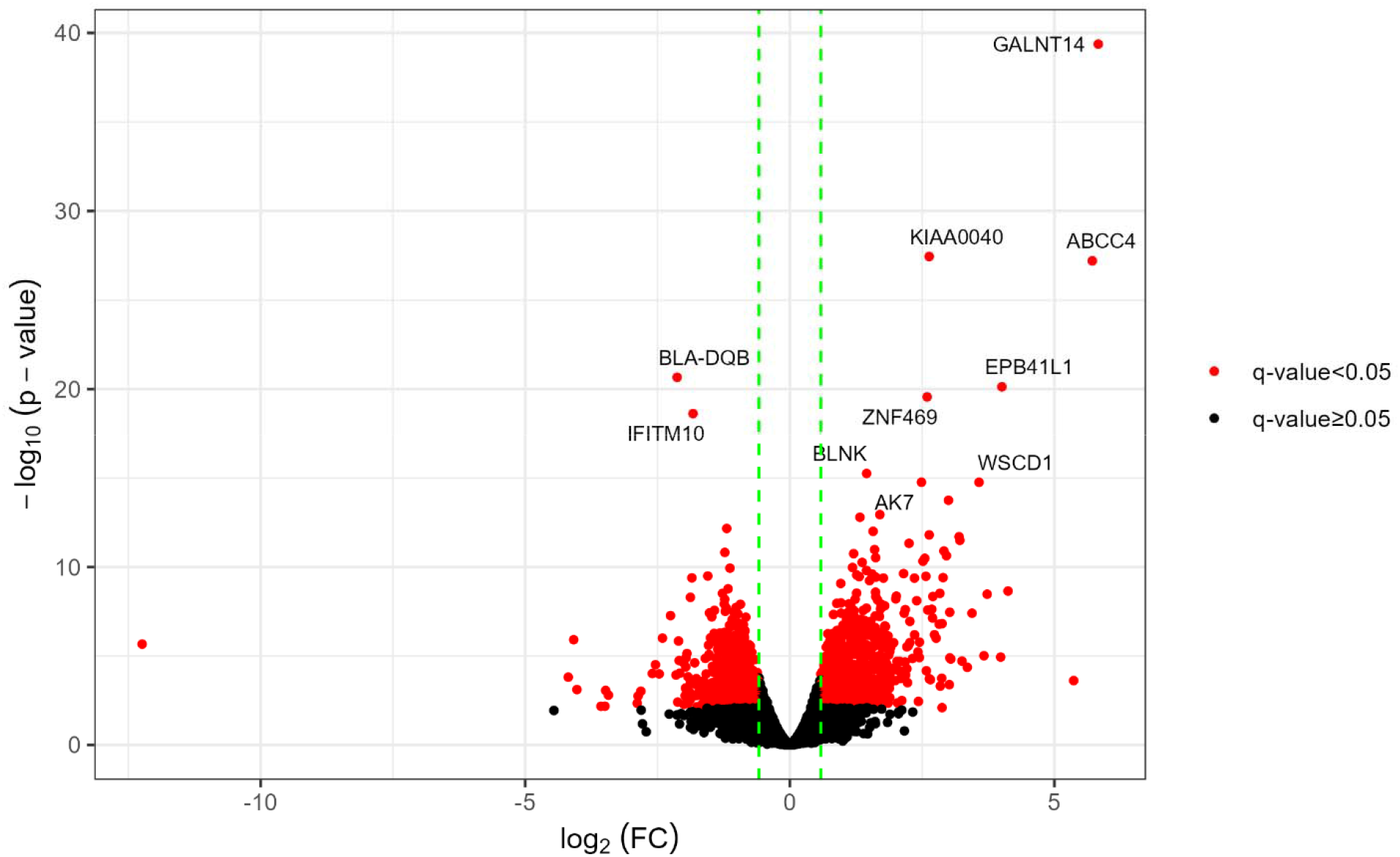
Main differentially expressed genes in HPVL vs LPVL transcriptomes. The vertical dotted green lines correspond to the log2 (FC) values of -1.5 and 1.5. The red dots denote DEGs and the black dots those statistically non-significant and/or with FC lower than the absolute value of 1.5 (|1.5|). Gene symbols indicate the 10 genes with the lowest q-values.

We then selected five genes showing a FC >1,5 and experimentally validated the differential expression in LPVL and HPVL cows via RT□qPCR. Four of these genes presented significant expression differences consistent with RNA-seq results (Figure 2), whereas *CD4* displayed a nonsignificant trend toward lower expression in HPVL animals.

**Figure 2.**
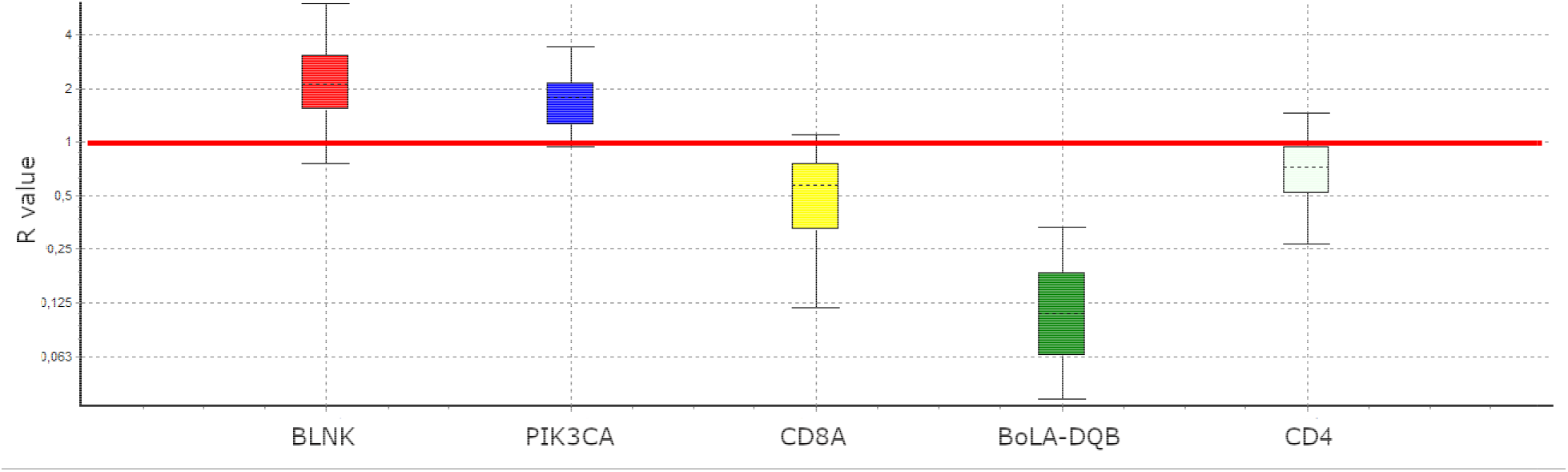
Experimental validation of differential gene expression by RT-qPCR expression ratio (R) for selected genes. The R were normalized against the geometric mean of the reference genes. *BLNK* (R = 2.192, p-value = 0.019) and *PIK3CA* (R = 1.728, p-value = 0.018) were UReg; conversely, *BoLA-DQB* (R = 0.492, p-value = 0.021) and *CD8A* (R = 0.108, p-value = 0.006) were DReg. CD4 (R = 0.7, p-value = 0.127) showed a tendency to DReg.

To gain functional insights into the phenotypes studied, we conducted gene enrichment analysis using the list of EGs ranked by logFC. The PANTHER and STRING databases were used to annotate Gene Ontology (GO) terms, Protein Class (PC) categories, and KEGG metabolic pathways, identifying enriched functional categories among upregulated (UReg) and downregulated (DReg) genes (Table S5). Several functional terms enriched in UReg genes were associated with cell division processes. The Biological Process (BP) GO-slim categories included “Mitotic cell cycle phase transition” (GO:0044772), “Protein localization to kinetochore” (GO:0034501), “DNA biosynthetic process” (GO:0071897), “Mitotic spindle assembly” (GO:0090307), and others (Table S5). Additionally, Protein Class (PC) terms enriched in UReg genes included “chromatin/chromatin-binding or -regulatory protein” (PC00077) and “Kinase activator” (PC00138), whereas KEGG pathway enrichment highlighted the “Cell cycle” (bta04110).

In contrast, most functional terms enriched in DReg genes were associated with immune responses, encompassing BP terms such as “Inflammatory response” (GO:0006954), “Response to interleukin-1” (GO:0070555), “Antigen processing and presentation” (GO:0019882), “T-cell-mediated immunity” (GO:0002456), and “Positive regulation of adaptive immune response” (GO:0002821). The significant KEGG pathways associated with the DReg genes included “Th1 and Th2 cell differentiation” (bta04658), “natural killer cell-mediated cytotoxicity” (bta04650), and “cytokine□cytokine receptor interaction” (bta04060). The enriched PC terms in the DReg genes were “Major histocompatibility complex proteins” (PC00149) and “Immunoglobulin receptor superfamily” (PC00124), both of which are key players in immune responses. Figure 3A and B show significant DEGs within the PC00149 and PC00124 terms, respectively.

**Figure 3.**
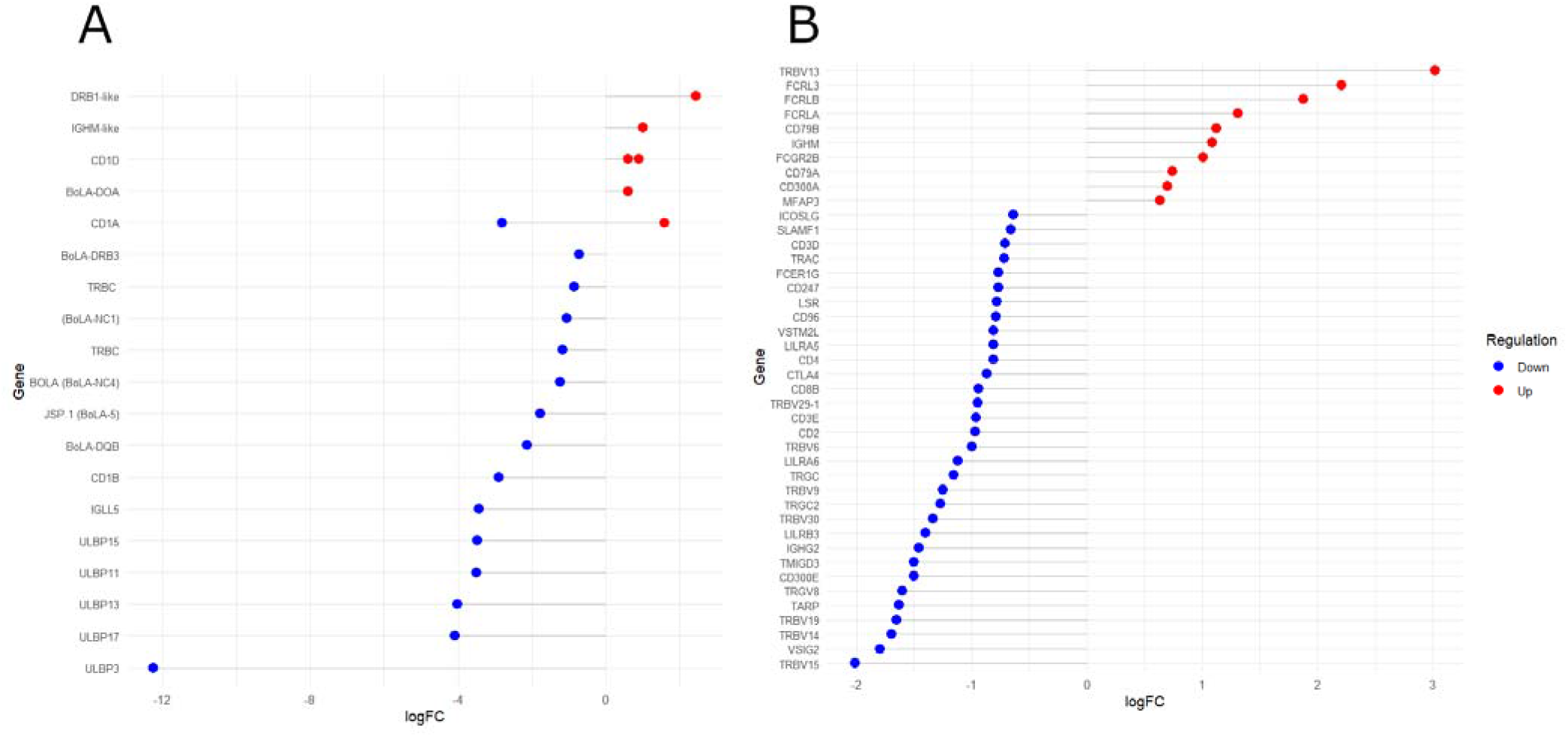
Lollipop chart of differentially expressed genes and their logFC values from two protein class (PC) PANTHER terms: A) “Major histocompatibility complex (MHC) proteins” (PC00149) and B) “Immunoglobulin (Ig) receptor superfamily” (PC00124). Red and blue dots indicate upregulated and downregulated genes, respectively, in HPVL animals.

A protein□protein interaction network based on the 1908 DEGs (FC ≥ |1.5|) consisted of 1732 nodes (proteins) and 10621 edges (interactions), forming a primary network (interaction score ≥ 0.4) with additional smaller subnetworks and isolated nodes (Figure S3). We selected the largest network to identify clusters of highly interconnected proteins, potentially representing protein complexes related to specific metabolic pathways. Six clusters were identified (Figure 4). Cluster A predominantly contained UReg genes, whereas clusters C, D, and E presented distinct subclusters of UReg (dotted circles) and DReg genes. Each cluster was analyzed for overrepresented GO, BP, and KEGG terms using the STRING database to interpret their biological functions (Table S6). Clusters enriched with UReg genes were associated with cell cycle regulation, mitotic division, and DNA repair, whereas clusters enriched with DReg genes were enriched in immunological processes, specifically innate and adaptive immunity. The functional terms and pathways enriched in the UReg and DReg genes were consistent across the enrichment analysis and cluster-based tests, validating the results through different analytical approaches. Notably, Cluster B (Figure 4) was enriched in terms related to sphingolipid metabolism.

**Figure 4.**
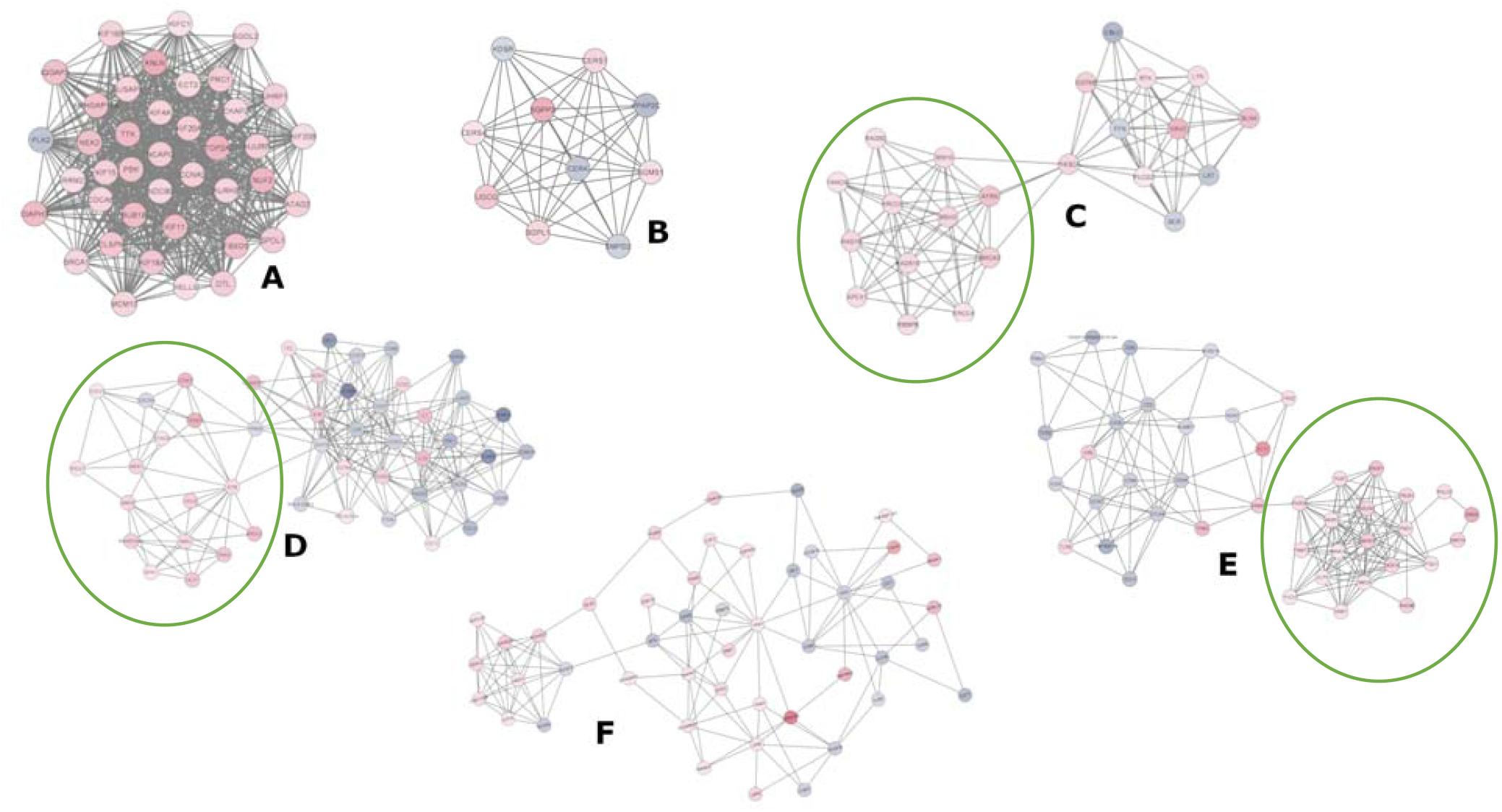
Clusters analysis of the protein-protein interaction (PPI) networks. The nodes represent proteins, and the edges (connecting lines) denote various types of interactions as suggested in the STRING db. Node colors are determined by the expression values (logFC); the blue scale represents DReg genes, while the red scale represents UReg genes. Clusters C, D, and E consisted of two distinct sub-clusters, comprising UReg (dotted line circle) and DReg genes.

## Discussion

To identify bovine genes and biological pathways underlying contrasting levels of BLV infection, we conducted whole-transcriptome analyses on PBMCs (peripheral blood mononuclear cells) obtained from BLV-infected Argentinian Holstein cows showing low and high BLV proviral load (LPVL and HPVL, respectively).

Differential expression analysis (HPVL vs. LPVL) revealed 2216 differentially expressed genes (DEGs) (q < 0.05) among the 13,382 genes evaluated. Among these DEGs, only 37.4% (829) presented high fold change values (FC ≥ |2|), whereas 86.1% (1,908) presented moderate changes (FC ≥ |1.5|). These findings agreed with previous reports of modest gene expression changes in blood cells in response to BLV infection (Brym and Kamiński, 2017).

Among the top 10 DEGs with the lowest q-values (Figure 1), eight were classified as UReg (upregulated), while two were categorized as DReg (downregulated). Notably, seven UReg genes (*GALNT14, KIAA0040, EPB41L1, ZNF469, BLNK, AK7*, and *WSCD1*) are associated with the progression of various cancers in humans (Peng et al., 2008; Xi et al., 2013; Tang and Zhang, 2018; Lin and Yeh, 2020; Tuluhong et al., 2020; Chacon-Barahona et al., 2021; Kurata et al., 2021). Conversely, two DReg genes, *BoLA-DQB* and *IFITM10*, are associated with immune processes. The *BoLA-DQB* gene encodes the β chain of the DQ molecule, which is essential for CD4+ T-cell activation and, hence, on the modulation of the adaptive cellular and humoral response (Norimine and Brown, 2005). *IFITM10*, a member of the interferon-induced transmembrane protein family, has been implicated in the inhibition of viral infections in humans, including HIV-1, Influenza, and SARS-CoV-1 (Almén et al., 2012).

The functional annotation revealed that immune-related categories, such as “antigen processing and presentation,” “positive regulation of adaptive immune response,” and “T-cell-mediated immunity,” predominantly included DReg genes (Table S5). A significant proportion of these immune-related genes were associated with the PANTHER term “Major histocompatibility complex (MHC) proteins” (PC00149). Figure 3A highlights the DEGs (q < 0.05, FC > |1.5|) within this category. Notably, *BoLA-DRB3* polymorphisms have been extensively investigated for their association with BLV proviral load (PVL) levels (Mirsky et al., 1998; Juliarena et al., 2008; Jimba et al., 2012; Miyasaka et al., 2013; Carignano et al., 2017; Takeshima et al., 2019). Additionally, other class II MHC genes (*BoLA-DQB, BoLA-DOA*, and *DRB1-like* pseudogene) were differentially expressed. These genes are expressed primarily in antigen-presenting cells (APCs), such as dendritic cells, mononuclear phagocytes, and B cells, where they play critical roles in regulating T-cell-mediated immune regulation. Variants in DRB3-DQB haplotypes have been associated with susceptibility or resistance to persistent lymphocytosis (PL) in BLV-infected cattle (Zanotti et al., 1996; Miyasaka et al., 2012; Fukunaga et al., 2020). Additionally, MHC class I (*BoLA-5*) and nonclassical MHC genes (*BoLA-NC1* and *NC4*) were downregulated in HPVL animals (Figure 3A). MHC class I molecules facilitate the presentation of intracellular viral peptides to CD8+ T cells (Hewitt, 2003), whereas nonclassical MHC genes regulate T-cell responses (Hofstetter et al., 2011). Dysregulation of CD1 class I-like genes, which present lipid antigens to T cells, was also observed. This finding is consistent with reports of altered CD1 expression in human viral infections (Chung et al., 2013; Shinya et al., 2016; Riou et al., 2017; Tan et al., 2018). Moreover, five DEGs encoding UL16-binding proteins (ULBPs), stress-induced molecules critical for NK-cell cytotoxicity, were significantly downregulated (Figure 3A). Downregulation of ULBPs has been linked to viral immune evasion mechanisms (Dunn et al., 2003; Jensen et al., 2011; Bauman et al., 2016).

Consistently, the PANTHER PC category “Immunoglobulin (Ig) receptor superfamily” (PC00124) was significantly enriched in DReg genes (Figure 3B). This category includes genes encoding T-cell receptors (TCRs) and components of the T-cell activation pathway. In contrast, the functional categories enriched in the UReg genes were predominantly related to the cell cycle, including “Mitotic cell cycle phase transition” (GO:0044772), “DNA biosynthetic process” (GO:0071897), and “assembly of the mitotic spindle” (GO:0090307). The KEGG pathway “Cell cycle” (bta04110) was also enriched in the UReg genes (Table S5). Dysregulation of these processes has been linked to oncogenesis, which is consistent with the oncogenic nature of BLV. Furthermore, the BP terms “B-cell activation” and “B-cell receptor signaling pathway” were overrepresented in the UReg genes. Approximately 30% of BLV-infected cows exhibit increased B-cell proliferation (Sordillo et al., 1994; Florins et al., 2008; Van den Broeke et al., 2010; Erskine et al., 2011), which has been associated with disrupted cell proliferation and apoptosis (Aida et al., 2013; Frie and Coussens, 2015).

The enrichment of terms such as “zinc-finger transcription factors (C2H2) (PC00248),” “acetyltransferases” (PC00042) and “histone acetylation” (GO:0016573) in UReg genes suggests potential epigenetic regulation. Notably, C2H2 zinc-finger proteins are involved in chromatin remodeling, a mechanism exploited by retroviruses to establish latency (Pluta et al., 2020). This may facilitate BLV immune evasion, promoting the expansion of infected B-cell clones.

Protein□protein interaction (PPI) network analysis (Figure S3) revealed clusters of highly interconnected proteins (Figure 4). The functional enrichment of clusters primarily composed of UReg genes revealed associations with cell division, mitotic spindle formation, and DNA repair (Clusters A, C-E; Figure 4; Table S6). In contrast, clusters predominantly composed of Dreg genes were enriched for immune response pathways, including T-cell metabolism and immune signaling (Clusters C-E, Figure 4, Table S6).

Cluster F, composed of both upregulated and downregulated genes in HPVL animals, was overrepresented in categories containing genes involved in cell cycle regulation and adaptive immune responses, respectively. In contrast, cluster B was enriched in terms related to sphingolipid metabolism. Cholesterol and sphingolipids form lipid rafts in the plasma membrane of B cells, which serve as platforms for increasing the concentration of B-cell receptors (BCRs) following antigen (Ag) stimulation (Aman and Ravichandran, 2000; Petrie et al., 2000). Notably, in BLV-infected animals, resistance to BCR translocation to lipid rafts has been observed (Hamilton et al., 2003). Additionally, sphingolipid metabolism plays critical roles in lymphocyte circulation and viral processes such as entry, replication, assembly, budding, and release, particularly in enveloped viruses (Yager and Konan, 2019; Schneider-Schaulies et al., 2021).

Overall, our findings suggest two nonmutually exclusive mechanisms contributing to the HPVL phenotype: dysregulation of B-cell proliferation/apoptosis and modulation of T-cell and NK-cell responses. Additionally, the suppression of proviral transcription via inhibitory transcription factors, such as zinc-finger proteins, may facilitate BLV immune evasion. Together, these mechanisms promote the expansion of infected B-cell clones, driving the HPVL phenotype. Given the increased risk of BLV transmission in HPVL cows (Gutiérrez et al., 2014a; Mekata et al., 2015; Juliarena et al., 2016), the segregation or elimination of HPVL animals from farms remains a viable control strategy (Ruggiero et al., 2019). However, middle/low-PVL animals in transition to an HPVL phenotype are present. In this study, we identified a reduced number of DEGs participating in protein□protein interaction clusters that could be further investigated as early biomarkers of an HPVL phenotype. Our the results are in line with findings previously reported by Nishimori et al. (2023) providing additional support. Indeed, five of the seven markers positively correlated with PVL (Nishimori et al., 2023) were found to be upregulated in HPVL animals in this work (Table S4).

Future studies should focus on the use of single-cell RNA-seq to uncover immune cell heterogeneity within the peripheral blood of infected cattle. This approach could help identifying subsets of B cells or T cells that may play a protective role against BLV proliferation in LPVL animals.

## Conclusion

The differential expression analysis conducted in this study identified a group of significantly differentially expressed genes (DEGs) when comparing animals with the HPVL and LPVL phenotypes in naturally infected cattle under productive conditions. The downregulated (DReg) genes were associated primarily with functions related to the cellular immune response, whereas the upregulated (UReg) genes were linked predominantly to the dysregulation of cell proliferation and the silencing of viral gene transcription. These mechanisms may enable the BLV to increase provirus copy numbers and evade the host’s immune response. This study enhances our understanding of the pathways involved in BLV infection and provides valuable insights that could contribute to the identification of infection markers, therapeutic targets, and information useful for BLV dissemination control programs within herds.

## Declarations

### Ethics approval

Not applicable.

### Authors’ contributions

- Conceptualization: MIP, MMM, KGT, HAC.
- Data curation: MIP.
- Formal analysis: MIP, HAC.
- Funding acquisition: MMM, KGT
- Investigation: MIP, GSA, HAC.
- Methodology: MIP, HAC.
- Supervision: MMM, KGT, HAC.
- Visualization: MIP.
- Writing – original draft: MIP, HAC.
- Writing – review & editing: MIP, GSA, MMM, KGT, HAC.
- Roles as defined by: CRediT (contributor role taxonomy)

### Availability of data and materials

The raw and processed data generated and analyzed in this study are available in the Gene Expression Omnibus (GEO) repository under the accession number GSE282244.

## Acknowledgments

We would like to thank Dr. Andrea Puebla and Pablo Vera from the Genomic Unit (IABIMO INTA-CONICET) for their assistance with library construction and next-generation sequencing.

We would also like to thank Dr. Irene Alvarez for allowing us to use laboratory resources for sample processing.

