## Supplementary material for "Whole-transcriptome analysis of BLV-infected cows reveals downregulation of immune response genes in high proviral loads cows": Figure S1, S2, S3 - Table S1, S2, S3

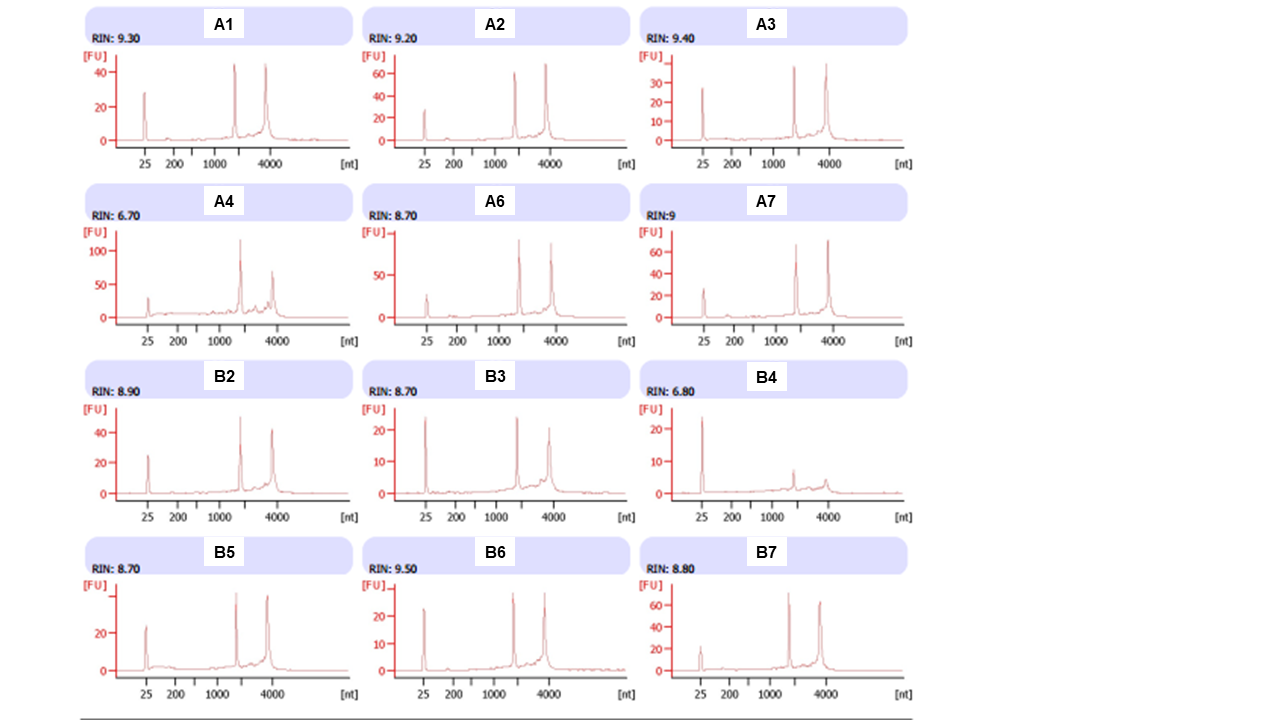

Figure S1: RNAs electropherograms. The fluorescence intensity [FU] versus the fragments size (nt) are showed. RIN: RNA integrity value.

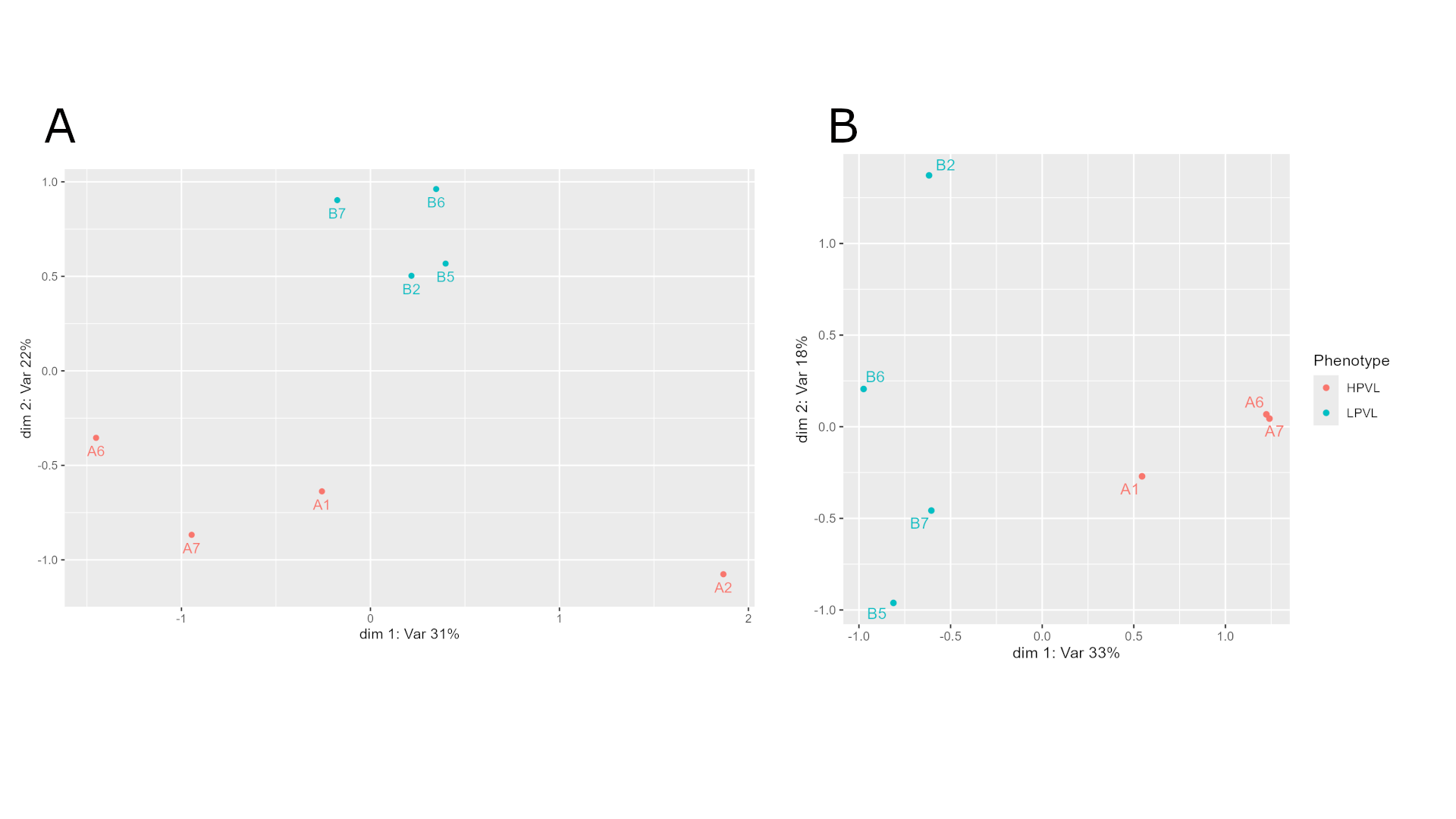

Figure S2: Multidimensional scaling (MDS) plots. Red dots depict high proviral load (HPVL) while blue dots represent the low proviral load (LPVL) animals. A) including the sample outlier A2 for the first dimension (dim 1), which explains the major variations. B) After exclusion of sample A2.

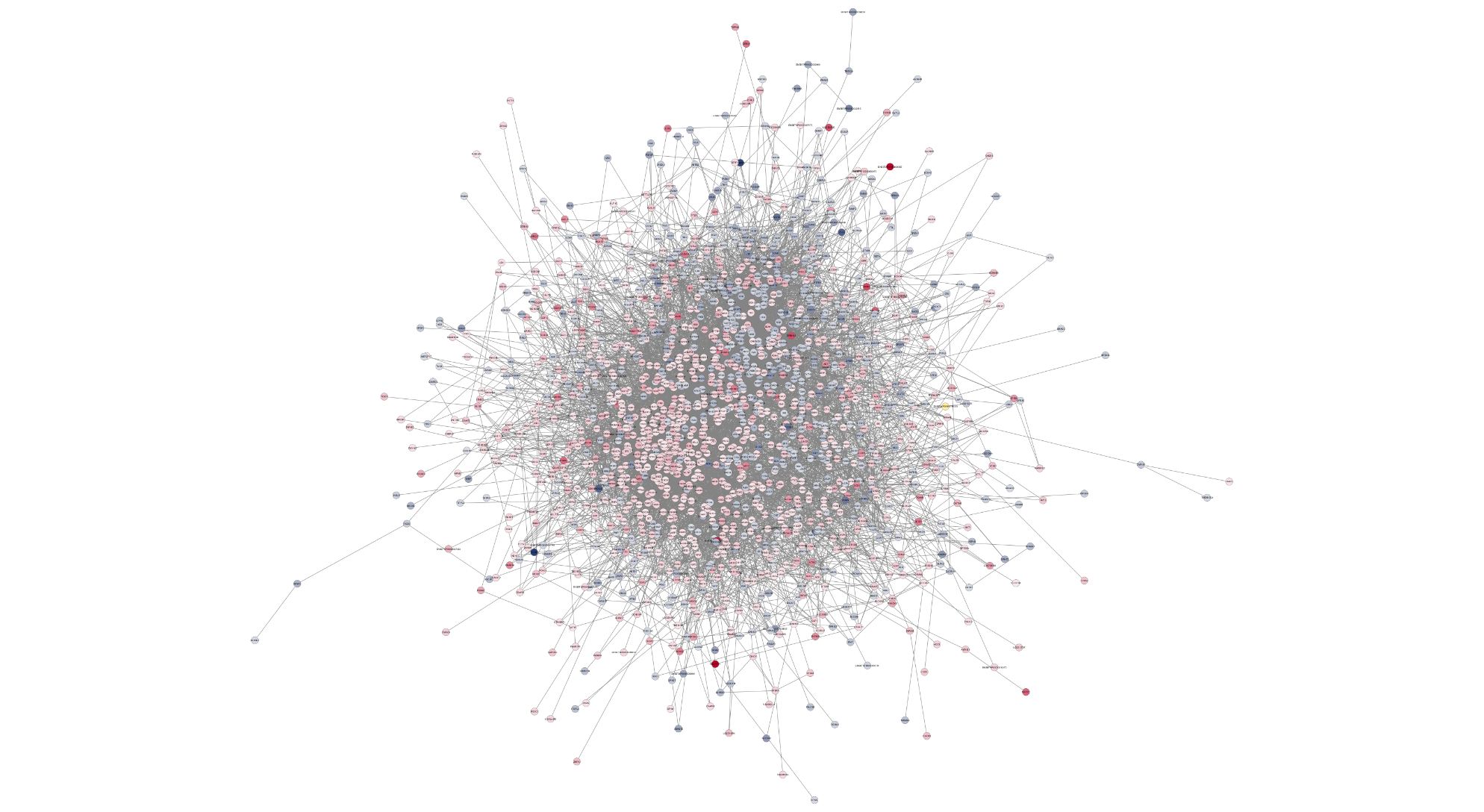

Figure S3: Protein-protein interaction (PPI) network representation of DEGs. The nodes represent proteins, and the edges (connecting lines) denote various types of interactions as suggested in the STRING db. Node colors are determined by the expression values (logFC); the blue scale represents DReg genes, while the red scale represents UReg genes.

Table S1: Summary of sequencing reads and mapping statistics per sample

| **Sample** | **A1** | **A2** | **A6** | **A7** | **B2** | **B5** | **B6** | **B7** |
| --- | --- | --- | --- | --- | --- | --- | --- | --- |
| **Reads number** | 49702459 | 43784088 | 32352588 | 46082521 | 47144198 | 50508228 | 41525184 | 43076731 |
| **Not mapped** | 1012469 (2.04%) | 892795 (2.04%) | 670340 (2.07%) | 817483 (1.77%) | 1091477 (2.32%) | 1487108 (2.94%) | 1493996 (3.60%) | 1263722 (2.93%) |
| **Uniquely mapped reads** | 43589906 (87.70%) | 37679909 (86.06%) | 28366477 (87.68%) | 39966339 (86.73%) | 40871104 (86.69%) | 43394830 (85.92%) | 35656883 (85.87%) | 37362801 (86.74%) |
| **Multi-mapped reads** | 5100084 (10.26%) | 5211384 (11.90%) | 3315771 (10.25%) | 5298699 (11.50%) | 5181617 (10.99%) | 5626290 (11.14%) | 4374305 (10.53%) | 4450208 (10.33%) |
| **Mapping rate** | 97.96% | 97.96% | 97.93% | 98.23% | 97.68% | 97.06% | 96.4% | 97.07% |
| **Mapped to DNA strand (+)** | 22236656 | 20610663 | 14171664 | 20581228 | 20884793 | 22703459 | 18431174 | 19278136 |
| **Mapped to DNA strand (-)** | 21353250 | 17069246 | 14194813 | 19385111 | 19986311 | 20691371 | 17225709 | 18084665 |
| **Mapped to exons (%)** | 33716461 (67.84%) | 30486086 (69.63%) | 22683071 (70.11%) | 31933183 (69.30%) | 32431605 (68.79%) | 35399305 (70.09%) | 28844793 (69.46%) | 30252305 (70.23%) |

Table S2: Candidate gene RT-qPCR reaction specifications.

| **Gene^a^** | **Primer sequence (5'→3')** | **Tm^c^** | **Homodimer (ΔG ^b^) ^ϒ^** | **Horquilla (Tm ^d^) ^ϒ^** | **Hairpin (ΔG^b^) ^ϒ^** | **Amplicon (pb)** | **Reference/Accession code^e^** | **log_2_FC^f^** |
| --- | --- | --- | --- | --- | --- | --- | --- | --- |
| *BLNK* | F-CGGATGACTTCGACAGCGATT | 57.3 °C | -6.76 Kcal/mol | 41.5 °C | 6.66 Kcal/mol | 188 | NM_001046054.2 | 1.45 |
|  | R-CCTCTGGCTTGATCGGTTGT | 57.4 °C | -4.62 Kcal/mol | -8.1 °C |  |  |  |  |
| *PIK3CA* | F-GGAGTCCTATTGCCGTGCAT | 57.5 °C | -7.05 Kcal/mol | 29.5 °C | -5.09 Kcal/mol | 168 | NM_174574.1 | 0.97 |
|  | R-AAATCTGGTCGCCGCATTTG | 56.6 °C | -5.36 kcal/mol | 3.5 °C |  |  |  |  |
| *BoLA-DQB* | F-CCTGTCAGCCTGTCCTACTC | 56.8 °C | -3.55 Kcal/mol | 10.4 °C | -5.13 Kcal/mol | 160 | NM_001034668.3 | -2.13 |
|  | R-GAAGCTCTTGGGGTCTGAGT | 56.6 °C | -6.34 Kcal/mol | 29.4 °C |  |  |  |  |
| *CD8A* | F-TGGACTTCGCCTGCAATATCT | 56.4 °C | -7.05 Kcal/mol | 24.4 °C | -7.04 Kcal/mol | 158 | NM_174015.1 | -0.92 |
|  | R-GTTGGGCTTGCCTCCTTGT | 58.4 °C | -6.21 Kcal/mol | 44.1 °C |  |  |  |  |
| *CD4* | F-AGCAGAAAGTGAAACTCGTGG | 55.4 °C | -3.61 Kcal/mol | 31.6 °C | -5.02 Kcal/mol | 148 | X. S. Wang et al., 2013 | -0.81 |
|  | R-ACCAACTTCGGCTGATTTGAG | 55.5 °C | -3.90 Kcal/mol | 24.2 °C |  |  |  |  |

^a^ GenBank gene symbol. ^ϒ^ Secondary structure parameters. ^b^ Gibbs free energy (ΔG) for the formation of homo and heterodimers (optimal > -9 Kcal/mol). ^c^ Melting temperature (Tm) (optimal: similar in both primers). ^d^ Tm for formation of primer secondary structure (optimal: lower than the primers pair). ^e^ GenBank accession code. ^f^ Log_2_FC: fold change values from the RNA-seq experiment.

Table S3: Samples proviral load at times T3 and T4.

| **Sample ID** | **T3** | **T4** | **Phenotype** |
| --- | --- | --- | --- |
| A1 | 30994.6 | 32081.7 | HPVL |
| A3 | 33660.5 | 54808 | HPVL |
| A5 | 79864.3 | 102647.5 | HPVL |
| A6 | 110437.3 | 119882.4 | HPVL |
| B2 | 173.6 | 166.1 | LPVL |
| B5 | 0 | 166.2 | LPVL |
| B6 | 0 | 192.5 | LPVL |
| B7 | 0 | 31.1 | LPVL |
| B8 | 0 | 0 | LPVL |

Table S4: Differentially expressed genes (DEGs) between HPVL and LPVL groups.

Table S5: Gene Ontology categories (BP, MF and CC), protein classes (PC) and KEGG metabolic pathways significantly enriched.

Table S6: Gene Ontology (GO) terms and KEGG metabolic pathways significantly overrepresented in clusters A-F.
